# A single CRISPR base editor to induce simultaneous C-to-T and A-to-G mutations

**DOI:** 10.1101/729269

**Authors:** Rina C. Sakata, Soh Ishiguro, Hideto Mori, Mamoru Tanaka, Motoaki Seki, Nanami Masuyama, Keiji Nishida, Hiroshi Nishimasu, Akihiko Kondo, Osamu Nureki, Masaru Tomita, Hiroyuki Aburatani, Nozomu Yachie

## Abstract

While several Cas9-derived base editors have been developed to induce either C-to-T or A-to-G mutation at target genomic sites, the possible genome editing space when using the current base editors remains limited. Here, we present a novel base editor, Target-ACE, which integrates the abilities of both of the previously developed C-to-T and A-to-G base editors by fusing an activation-induced cytidine deaminase (AID) and an engineered tRNA adenosine deaminase (TadA) to a catalytically impaired *Streptococcus pyogenes* Cas9. In mammalian cells, Target-ACE enabled heterologous editing of multiple bases in a small sequence window of target sites with increased efficiency compared with a mixture of two relevant base editor enzymes, each of which may block the same target DNA molecule from the other. Furthermore, by modeling editing patterns using deep sequencing data, the editing spectra of Target-ACE and other base editors were simulated across the human genome, demonstrating the highest potency of Target-ACE to edit amino acid coding patterns. Taking these findings together, Target-ACE is a new tool that broadens the capabilities for base editing for various applications.

---

CRISPR–Cas9 is a genome editing tool in which Cas9 is recruited by a guide RNA (gRNA) to its target DNA region containing a protospacer adjacent motif (PAM) in its 3′ end to induce a double-strand DNA break (DSB)^1,2^. This method has rapidly expanded our ability to knock out genes through error-prone DNA repair and to insert transgenes into chromosomes through DSB-induced homologous recombination (HR). Moreover, the DSB-induced HR also enables the substitution of targeted single bases by replacing a genomic sequence with a donor DNA template^3^ for applications ranging from functional testing of genetic traits to developing clinical methods to correct disease mutations. However, this strategy for base substitutions through DSB has been shown to cause cell toxicity^4^ and chromosomal deletions and translocations^5^ and the low HR efficiency makes the introduction of multiple substitutions in different loci difficult.

In contrast, Cas9-derived base editors employ tethering of a deoxynucleoside deaminase to a nuclease-deficient or nickase Cas9 (dCas9 or nCas9)–gRNA complex to induce efficient and direct base substitutions in the genomic sequence^6^. Among the currently available base editors, cytosine base editors (CBEs)^7,8^ and adenine base editors (ABEs)^9^ enable highly efficient and precise base substitutions in a narrow window of gRNA targeting sites. CBEs commonly consist of two components fused to dCas9 or nCas9: a cytidine deaminase, such as rAPOBEC1 used in BEs^8,10^ and PmCDA1 used in Target-AID^7^, to convert cytidines of the non-target strand into uridines by deamination; and a uracil glycosylase inhibitor (UGI) that inhibits base excision repair, allowing uracil bases to be replaced with thymine bases through the mismatch repair pathway and DNA replication (C-to-T base editing). ABEs instead utilize adenosine deaminases that convert adenosines to inosines^9^. While none of the naturally existing enzymes is known to catalyze the deamination of deoxyadenosines, directed protein evolution of an *Escherichia coli* tRNA adenosine deaminase (TadA) has been carried out and a heterodimer complex of wild-type and mutant TadA was found to exhibit the activity of converting targeted adenosines to inosines when fused to dCas9 or nCas9. As inosine bases are replaced with guanine bases through DNA replication, ABEs enable A-to-G base editing. The base editing efficiencies of CBEs and ABEs are elevated when nCas9 (D10A) is used to nick the target DNA strand complementary to the gRNA as the other strand with mutated bases is used as a repair template in the DNA nick repair process.

These base editors hold great promise in therapeutics as they can efficiently induce targeted base substitutions in chromosomes with low indel frequencies and minimal cell toxicity. While the currently available base editors enable four different transition mutations, C-to-T, G-to-A, A-to-G, and T-to-C, some of which have been demonstrated to correct or generate disease mutations in animal models^11,12^, there is no base editor that enables any of the other patterns. While no practical solution to develop base editors for the other eight non-deamination-mediated transversion mutations has yet been reported, a single base editor with both C-to-T and A-to-G base substitution activities would broaden the capability for genome editing, such as alteration of amino acid coding patterns. To this end, we have developed a new base editor, Target-ACE (adenine and cytosine editor), which induces simultaneous C-to-T and A-to-G base substitutions.

## Dual functionality of Target-ACE

We fused PmCDA1 of Target-AID and a heterodimer of TadA of ABE-7.10 with nCas9 (D10A) to develop Target-ACE (Fig. 1a). Previous studies reported that the C-terminus fusion of TadA and rAPOBEC1 abolished the base editing activity^9^, whereas efficient base editing was preserved in Target-AID fusing PmCDA1 to the C-terminus end^8,13^. Thus, we fused the TadA domain and PmCDA1 to the N- and C-termini of nCas9, respectively. Target-ACE was also designed to have a nuclear localization signal (NLS) and a C-terminal UGI, which increases the purity of C-to-T substitution.

**Figure 1.**
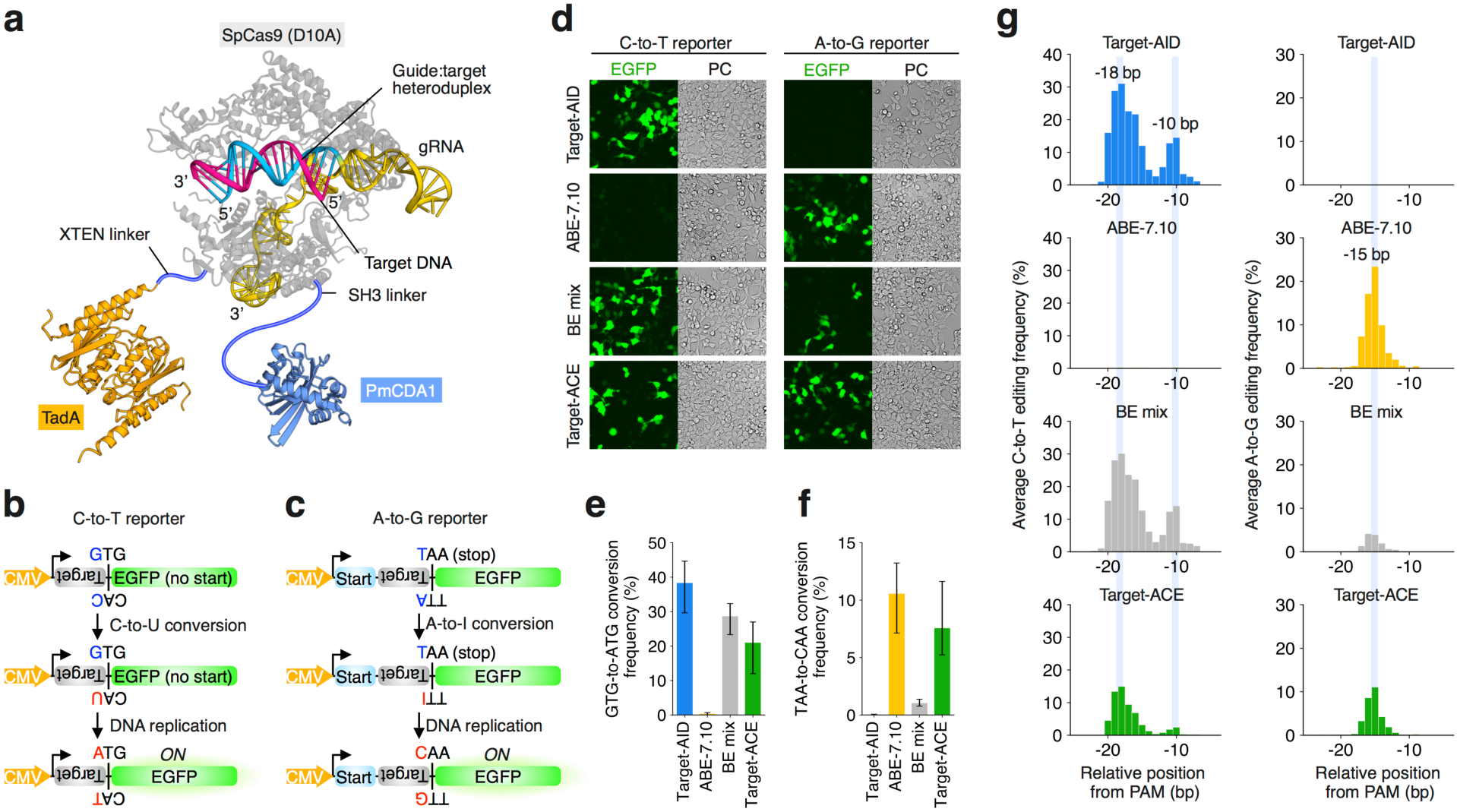
Target-ACE. (**a**) Structural overview of Target-ACE. The TadA heterodimer (orange) and PmCDA1 (light blue) are fused to the N-terminus and C-terminus ends of nCas9 (D10A) (light gray) through XTEN and SH3 linkers, respectively. NLS and UGI are not displayed in this diagram. (**b**) Schematic representation of the C-to-T base editing reporter. C-to-T base editing of the gRNA target strand followed by DNA replication restores the translation of EGFP by converting a mutated start codon GTG (valine) to ATG. (**c**) Schematic representation of A-to-G base editing reporter. A-to-G base editing of the gRNA target strand followed by DNA replication breaks a stop codon TAA to CAA (guanine) and releases the translation of its downstream EGFP. (**d**) Transfection of different base editors (or their combination) to cell lines harboring C-to-T or A-to-G base editing reporters with their corresponding gRNAs. (**e**) Average frequencies of the start codon restoration in the C-to-T editing reporter cells by different base editor reagents. Error bars show maximum and minimum frequencies of triplicates. (**f**) Average frequencies of the stop codon destruction in the A-to-G editing reporter cells by different base editor reagents. (**g**) Average C-to-T and A-to-G base editing spectra of different base editor reagents for endogenous targeting sites in HEK293Ta cells.

To test the base editing activity of Target-ACE and other base editors in living cells without sequencing, we first constructed C-to-T and A-to-G base editing reporter circuits, in which the corresponding base substitutions were designed to activate EGFP protein translation. The C-to-T reporter circuit was designed to restore a mutated GTG start codon to ATG by C-to-T base editing of its antisense strand (Fig. 1b). In the A-to-G reporter, EGFP translation is released when a TAA stop codon encoded in its downstream region is broken to CAA by A-to-G base editing in its antisense strand (Fig. 1c). The C-to-T and A-to-G reporter circuits were introduced into human embryonic kidney HEK293Ta cells by lentiviral transduction. To each of the reporter cell lines, we then transfected base editor expression vectors with the reporter targeting gRNAs and observed their fluorescence activities. From the C-to-T reporter cells, EGFP expression was observed for Target-AID, Target-ACE, and the pooled transfection of Target-AID and ABE-7.10 expression vectors (BE mix), but not for ABE-7.10 (Fig. 1d and Supplementary Fig. 1). From the A-to-G reporter cells, EGFP expression was observed for ABE-7.10 and Target-ACE, but not for Target-AID, whereas slightly reduced EGFP induction was detected for BE mix (Fig. 1d and Supplementary Fig. 1).

Using amplicon sequencing, we then quantified the editing frequencies of the different base editing methods at the target regions for the reporter cell samples. The GTG-to-ATG conversion frequencies in the C-to-T reporter cell samples for Target-AID, ABE-7.10, BE mix, and Target-ACE were 38.43%, 0.23%, 28.56%, and 20.80%, respectively (Fig. 1e). The TAA-to-CAA conversion frequencies in the A-to-G reporter cell samples for Target-AID, ABE-7.10, BE mix, and Target-ACE were 0.04%, 10.60%, 1.03%, and 7.57%, respectively (Fig. 1f). Target-ACE showed base editing activities for both GTG-to-ATG and TAA-to-CAA conversions, which were, however, mildly diminished compared with the GTG-to-ATG conversion efficiency of Target-AID (fold change of 0.54) and the TAA-to-CAA conversion efficiency of ABE-7.10 (fold change of 0.71). Notably, consistent with the fluorescence imaging experiment, the TAA-to-CAA conversion efficiency of ABE-7.10 was largely diminished when it was mixed with Target-AID. Given that the Cas9–gRNA complex remains bound to the target DNA molecule even after editing^14^, this suggests that Target-AID prevents the access of ABE-7.10 to the same target DNA molecule.

## Base editing spectrum of Target-ACE for targeting endogenous regions

To test editing spectra of different base editing methods, we prepared 23 gRNAs targeting human chromosomal regions (Supplementary Table 1). Eight gRNAs were designed to target regions having poly-cytosine (poly-C) sequences in various positions relative to the PAM, while another set of eight gRNAs were similarly designed to target poly-adenine (poly-A) regions. The other seven gRNAs were designed to target alternating adenine and cytosine repeats; three had adenines and cytosines at even and odd positions, respectively, relative to the PAM (poly-AC), while four had the opposite arrangement (poly-CA). Each of the gRNA expression vectors was transfected into HEK293Ta cells with different base editor reagents and the base editing patterns were analyzed by amplicon sequencing of the target region (Supplementary Figs. 2–4).

The editing efficiencies of different base editing methods for the chromosomal regions were consistent with those observed in the C-to-T and A-to-G reporter cell assays. Among the poly-C, poly-AC, and poly-CA targets, the average frequencies of amplicon sequencing reads with C-to-T substitutions were on par for Target-AID and BE mix (33.84% and 34.72%, respectively), while that for Target-ACE was reduced (15.24%) (Supplementary Fig. 5). In contrast, among the poly-A, poly-AC, and poly-CA targets, the average frequencies of reads with A-to-G substitutions induced by ABE-7.10 (23.80%) were largely diminished upon mixing with Target-AID (5.20%), while Target-ACE exhibited A-to-G base editing activity with an alleviated functional deficiency (12.26%) (Supplementary Fig. 6).

By taking the average of C-to-T and A-to-G base editing frequencies for every target sequence position relative to PAM, we also confirmed that the C-to-T and A-to-G editing spectra of Target-ACE were similar to those of Target-AID and ABE-7.10, respectively (Fig. 1g). Although we previously reported that the C-to-T base editing activity of Target-AID was in a restricted area of a target region with the peak activity position at −18 bp from PAM^15^, we found in this study that the editable region of Target-AID for C-to-T base editing with ≥1% frequency spanned from −20 bp through −7 bp with two activity peaks at the positions of −18 bp as well as −10 bp (with peak heights of 31.00% and 14.45%, respectively). ABE-7.10 showed an A-to-G editable region spanning from −17 bp through −12 bp with an activity peak at −15 bp (23.37%), consistent with a previous report^9^. For Target-ACE, the editable region spanned from −20 bp to −10 bp for C-to-T editing, with activity peaks at −18 bp (14.90%) and −10 bp (2.41%) and from −17 bp to −13 bp for A-to-G editing with the activity peak at −15 bp (11.02%). Taken together, these results demonstrate that Target-ACE enables the C-to-T and A-to-G editing at target regions, whereas the simultaneous use of Target-AID and ABE-7.10 efficiently mediates the C-to-T, but not A-to-G, editing.

## Simultaneous editing of heterologous nucleotide bases

Multiple cytosines of the poly-C regions and adenines of the poly-A regions were frequently co-edited by Target-AID and ABE-7.10, respectively (Supplementary Figs. 7 and 8). We also found that simultaneous C-to-T editing and A-to-G editing were induced in the seven poly-AC and poly-CA regions by Target-ACE and BE mix. Furthermore, the co-editing frequencies of Target-ACE were markedly higher than those of BE mix for 31 out of 33 cytosine–adenine pairs, which showed co-editing frequencies of ≥1% by either of the two base editing methods (Fig. 2a). Furthermore, consistent with the editing spectra of Target-ACE for C-to-T and A-to-G base substitutions, the co-editing frequency of heterologous bases was generally higher, for which cytosines and adenines were closer to the positions −18 bp and −15 bp relative to PAM, respectively. Indeed, the highest co-editing frequency was observed for the –18C:–15A pair among the entire heterologous cytosine–adenine pairs in all of the three poly-CA regions involving the –18C:–15A pair. The co-editing frequencies of heterologous bases by Target-ACE ranged from 0.89- to 5.87-fold compared with those by BE mix, with average fold difference of 2.53; the larger differences were observed for the cytosine–adenine pairs for which A-to-G substitution frequencies were markedly lower for BE mix (Fig. 2b and Supplementary Fig. 6).

**Figure 2.**
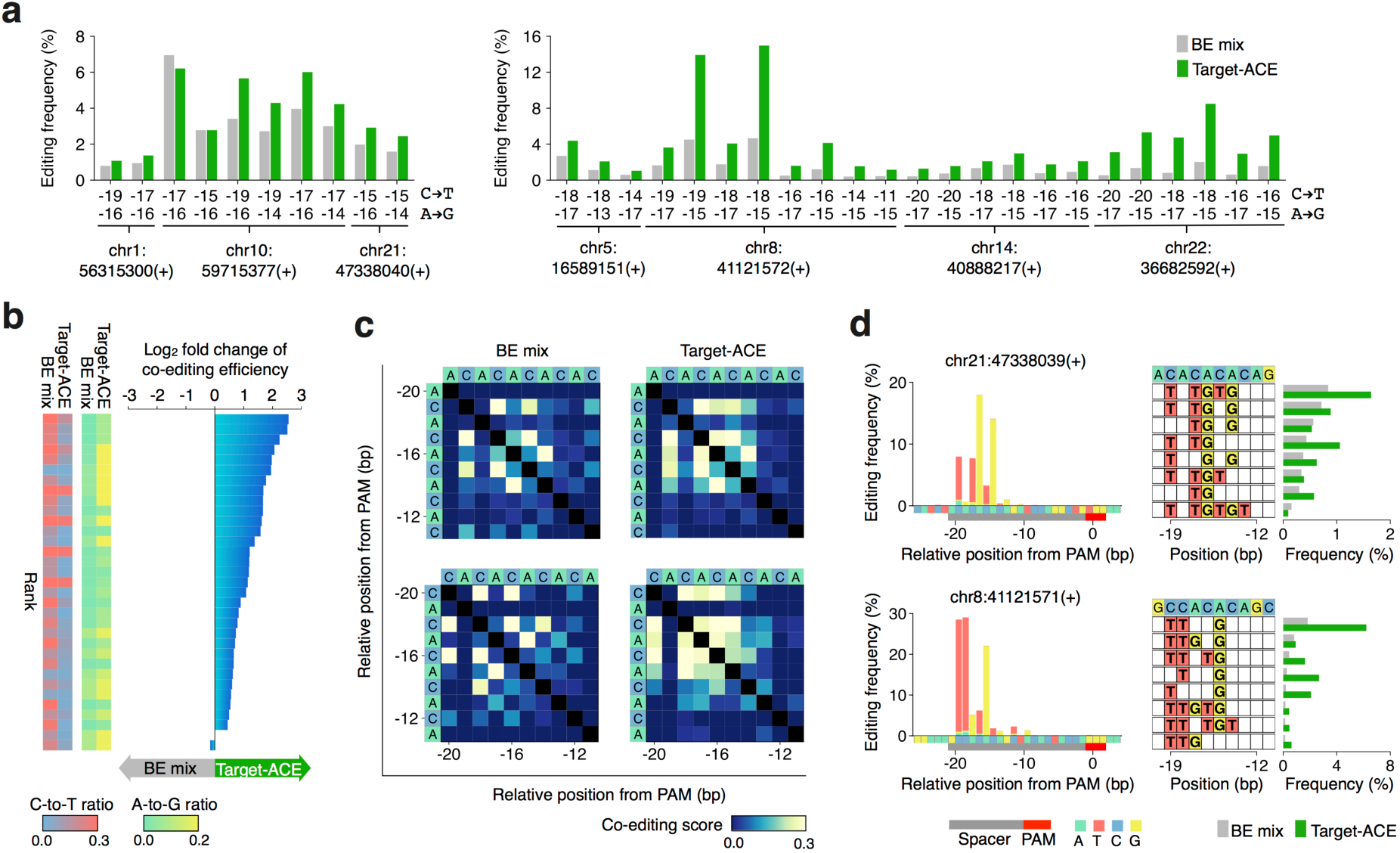
Simultaneous C-to-T and A-to-G base editing induced by Target-ACE and BE mix. (**a**) Frequencies of simultaneous base editing of cytidines and adenines at different combinatorial positions relative to PAM in poly-AC and poly-CA target regions. Each target region is denoted by its chromosome and the 5′ end position and strand direction of PAM. (**b**) Comparison of simultaneous base editing frequencies by Target-ACE and BE mix for the same heterologous base combinations. For each base combination, the fold change in efficiency of Target-ACE to BE mix is represented along with the independent C-to-T and A-to-G editing frequencies induced by these methods. (**c**) Average co-editing spectra of Target-ACE and BE mix for cytidines and adenines in the region from −20 bp to −11 bp relative to PAM of poly-AC and poly-CA sequences. (**d**) Base editing spectra for Target-ACE and frequencies of multi-base editing outcomes induced by Target-ACE and BE mix for a poly-AC region and a poly-CA region arbitrarily selected. In the left panels, base editing patterns and frequencies at different positions are represented by the color-coded x-axis for source nucleotide bases and the color-coded bars for destination nucleotide bases with their frequencies. In the right panels, the multi-base editing outcomes are shown for the top eight most common outcomes preferentially induced by BE mix.

To explore the simultaneous editing spectra of the different base editing methods, we calculated their co-editing scores for all pairs of two cytosine and/or adenine nucleotides in the poly-AC and poly-CA regions, where the co-editing score refers to the intersection-over-union for substituted bases in two different positions (Supplementary Figs. 9–12). While Target-AID and ABE-7.10 showed only co-editing activities against the same transition mutations and the average co-editing profiles of BE mix with the minimal A-to-G editing activity were similar to those of Target-AID, Target-ACE showed high co-editing activity for both adenines and cytosines in a restricted area relative to the PAM position (Fig. 2c and Supplementary Fig. 13). We also examined the efficiencies of more complex base editing patterns comprising both C-to-T and A-to-G substitutions for more than two positions by Target-ACE and BE mix. In five of the seven poly-AC and poly-CA target regions, Target-ACE outperformed BE mix in more than half of the top eight outcomes that BE mix preferentially produced (Fig. 2d and Supplementary Fig. 14). This result also demonstrated that the simultaneous C-to-T and A-to-G base editing activity of Target-ACE is higher than that of BE mix.

## Computational modeling of base editing outcomes

While recent studies reported that wild-type Cas9-mediated genome-editing outcomes can be predicted by machine learning approaches from sequences of target regions^16-18^, there have been no approaches to computationally predict base-editing outcomes. We thus sought to establish a base-editing prediction framework by simple modeling of conditional base transition probabilities (Fig. 3a). Using this model, we predicted the frequencies of all of the observed editing outcomes in each of the 23 target regions by training the other target regions, and found that the predicted frequencies were significantly correlated with the actual measurements for all of the base editing methods (*P*-value < 10^−24^) (Fig. 3b and Supplementary Fig. 15).

**Figure 3.**
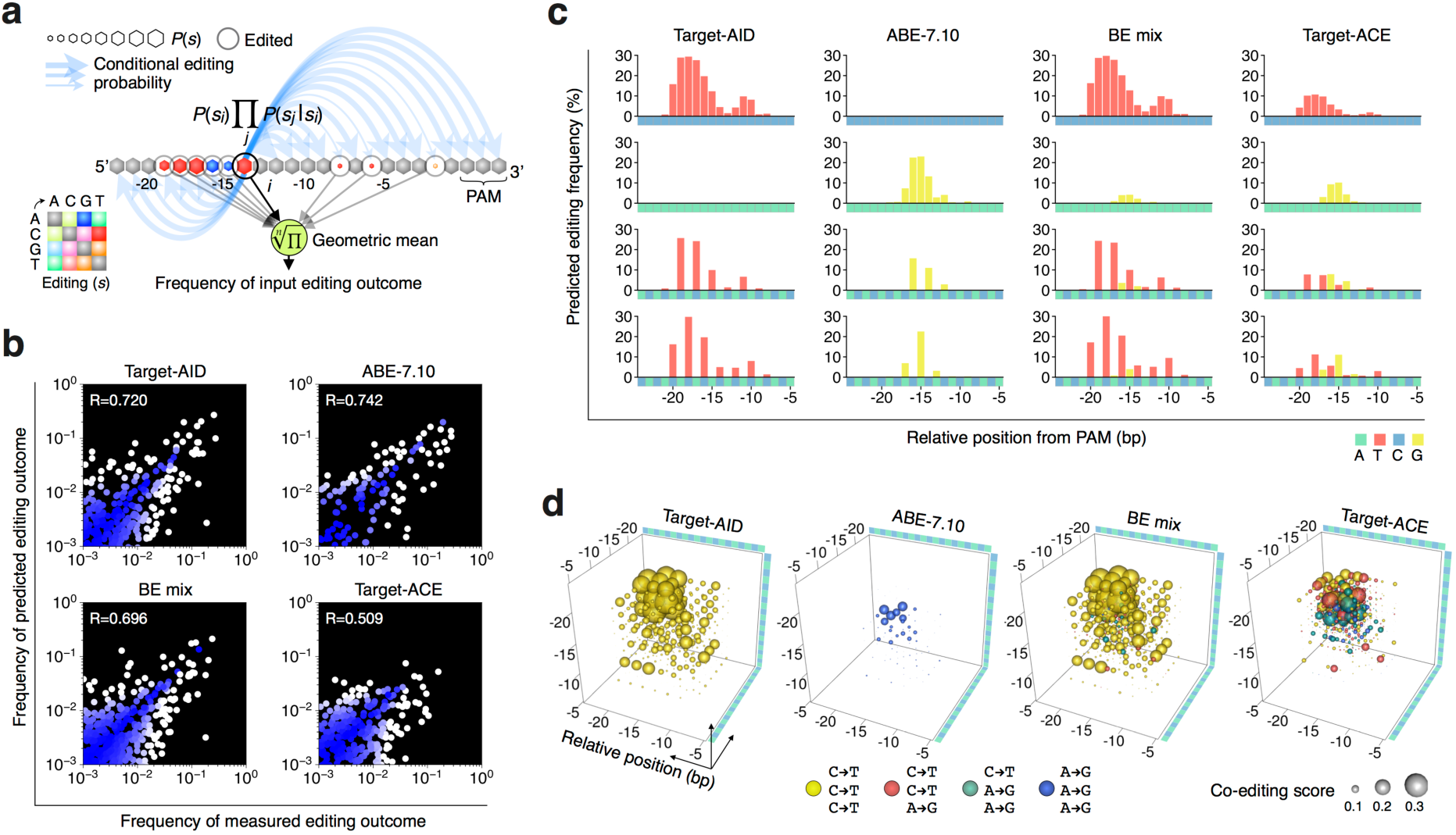
A prediction framework to simulate editing patterns induced by base editors. (**a**) Schematic diagram of the probabilistic model used in this study to predict frequencies of base editing outcomes. Briefly, to train a given base editor model, from a training amplicon sequencing data set, probabilities of single base editing events and their conditional probabilities given the other events are calculated thoroughly for different positions relative to PAM. The frequency of a given editing outcome in a test region is predicted as a geometric mean of a product of a probability of single editing status *s*_*i*_ with base transition and conditional probabilities of editing statuses in all the other positions given *s*_*i*_. (**b**) Correlation of measured and predicted editing outcome frequencies observed by leave-one-out cross-validation for each base editing method. The blue–white color scale indicates the Euclidean distance from the perfect prediction (diagonal). (**c**) Simulated base editing spectra of different base editing methods for poly-C, poly-A, poly-AC, and poly-CA regions continuously spanning from −24 bp through −5 bp from PAM. (**d**) Simulated co-editing spectra of different base editing methods for editing three nucleotide base positions in a poly-CA sequence.

To abstract the multi-dimensional editing spectrum of each base editing method, we next predicted the frequencies of single and combinatorial base conversion patterns induced in simulated poly-C, poly-A, poly-AC, and poly-CA sequences spanning from −24 bp through −5 bp relative to PAM. The single base editing spectra (Fig. 3c) and co-editing score profiles (Supplementary Fig. 15) predicted for different base editing methods were similar to those obtained from the experimental assays, also supporting the validity of the prediction framework. Furthermore, while the sequencing read depth limited the evaluation of tri- or more-base conversion frequencies from the actual amplicon sequencing datasets, the trained models allowed us to predict tri-nucleotide co-editing score profiles for different base editing methods (Fig. 3d). In this prediction, we found that the heterologous tri-nucleotide editing scores were almost always reduced for BE mix, whereas Target-ACE exhibited similar scores for both heterologous and homologous tri-nucleotide editing patterns.

## Codon conversion potentials of different base editing methods

Finally, we examined whether the simultaneous C-to-T and A-to-G base editing activity of Target-ACE expands the potential editing space of protein-coding patterns in the human genome. For each of 11,250,496 source codons in the human genome (corresponding to 20,062 RefSeq genes), we first predicted the frequencies of codon conversions to different destination codons by all of the possible gRNAs under the different base editing models, where each codon conversion was scored to be successful when the surrounding ±15 bp region was intact. The potential of conversion of each codon to a given destination codon was then defined to be its maximum predicted conversion frequency among those by the possible gRNAs for the same genomic position. Finally, we generated a codon convertibility profile (CCP) for each of the different base editing methods, where each source–destination codon pair was scored by the fraction of the source codons in the genome with conversion potential exceeding or equal to a given threshold (Fig. 4a and Supplementary Figure 16). As previously reported^19^, the potential codon conversion spaces of Target-AID and ABE-7.10 were both limited across all of the source–destination codon pairs, and their CCPs showed almost mirror images largely because of the symmetricity between the C•G-to-T•A and A•T-to-G•C base transitions. The CCP of BE mix was also similar to that of Target-AID as expected from the similarities in base editing patterns observed in the other analyses in this study. When the conversion profile threshold of 3% was applied, Target-ACE showed 24 more codon conversion patterns corresponding to 18 amino acid conversions compared with the union of the other base editing methods. In contrast, the numbers of targetable codons with conversion potential of ≥3% for different destination codons were reduced to 0.40-fold on average from Target-AID to Target-ACE for those targetable by Target-AID. A similar reduction in number of targetable codons was also observed for the equivalent comparison with ABE-7.10 (0.31-fold). The same values of fold change for BE mix were 1.00 and 0.05 compared with Target-AID and ABE-7.10, respectively. Nevertheless, out of the 60 codon conversion patterns that require both C•G-to-T•A and A•T-to-G•C base transitions, 50 patterns (48 non-synonymous substitutions; 30 amino acid substitutions) showed significantly higher codon conversion potentials for Target-ACE than for BE mix (Fig. 4b).

**Figure 4.**
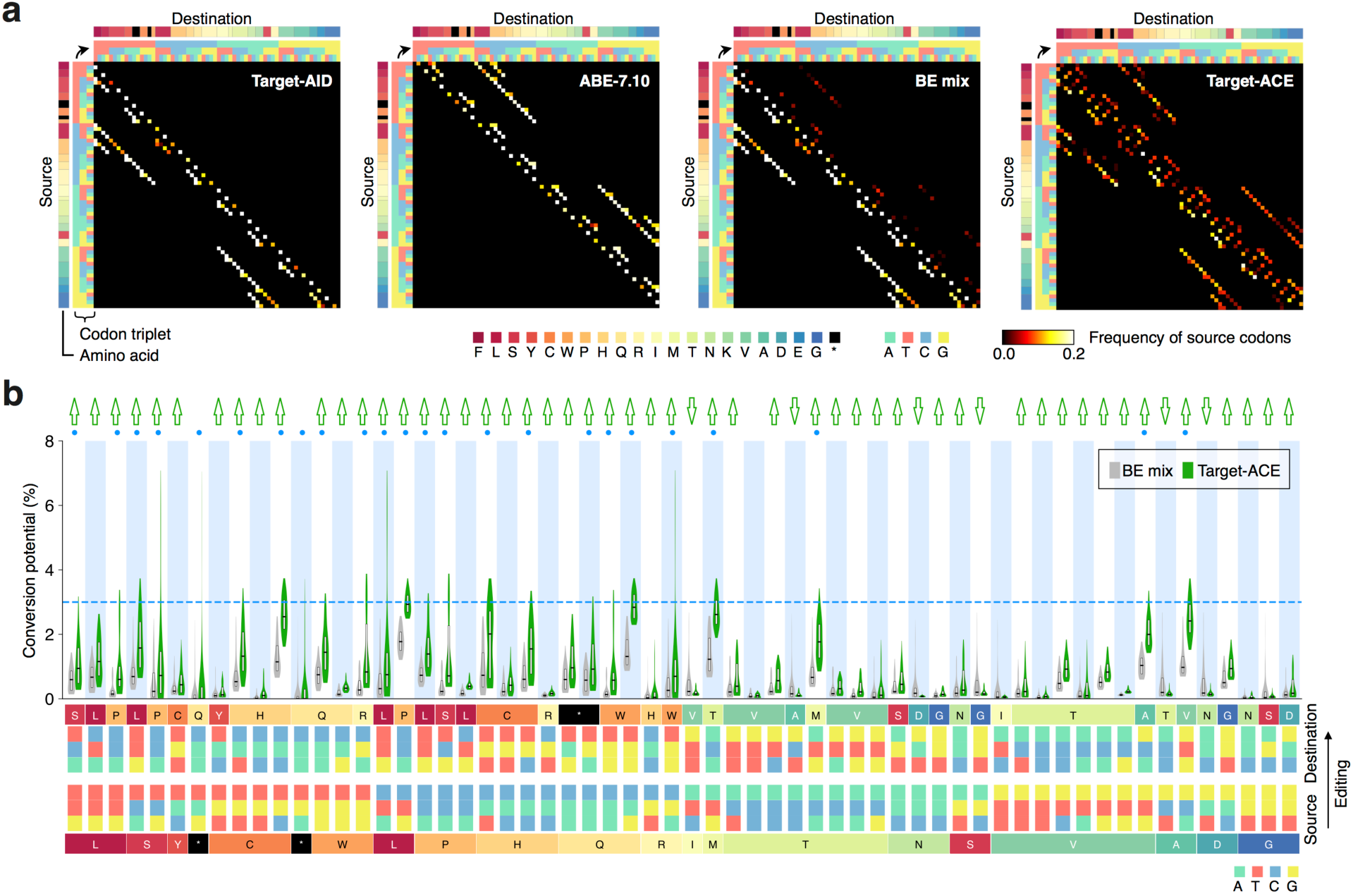
Simulation of genomic codon conversions by different base editing methods. (**a**) Codon convertibility profiles (CCPs) of different methods. For each source-to-destination codon conversion, the color scale indicates the fraction of the source codons in the human genome with predicted codon conversion potential of ≥3% that keep surrounding nucleotide regions intact. (**b**) Predicted codon conversion potential distributions of Target-ACE and BE mix for source–destination codon pair patterns each of which requires both C-to-T and A-to-G base substitutions. Green up and down arrows indicate codon pair patterns for which the conversion potential distribution for Target-ACE was significantly higher and lower than for BE mix, respectively (P-value < 10^−20^). Blue dots represent codon pair patterns with detected conversion potential of ≥3% (indicated by the blue dashed line).

## Discussion

While a recent study has reported a RNA base editor that utilizes dCas13 fused to an engineered ADAR2 retaining both the wild-type A-to-I and engineered C-to-U editing activities^20^, Target-ACE is the first that has demonstrated simultaneous C-to-T and A-to-G base substitutions in a narrow window of a target DNA region in a highly efficient manner. The simultaneous editing of heterologous bases by BE mix was inefficient, since the ABE-7.10-mediated A-to-G editing was largely abolished at most of the target sites. This suggests that Target-AID preferentially binds to a target DNA molecule through a gRNA and prevents the access of ABE-7.10, whereas the C-to-T and A-to-G activities of Target-ACE are free from this competition for access to targets. As shown in our simulation analyses, Target-ACE expands the potential space for editing amino acid coding patterns in the genome compared with the other CBEs and ABEs, highlighting its potential for therapeutic applications including the correction of disease-related mutations. Target-ACE could also be applied for *in vivo* mutagenesis. CRISPR-X involving a mutant human AID has been developed as a sequence diversification tool to induce C-to-A/G/T base substitutions for around a hundred base pair region of a target site, albeit with low efficiency at each position^21^. TAM incorporating another mutant human AID enables C-to-A/G/T base substitutions with high efficiency, but for a narrow editing window^22^. Although Target-ACE is also limited in the width of its editable region, this could somehow be supplemented by the use of multiple gRNAs, as previously demonstrated using TAM^22^. The strategy of multi-domain fusion employed in Target-ACE would at least drive the development of other base sequence diversification tools with broader base transition ability.

The estimated potential of precise codon conversion by Target-ACE was similar for all source–destination codon pairs that require C•G-to-T•A and/or A•T-to-G•C editing, whereas Target-AID and ABE-7.10 were suggested to surpass Target-ACE in conversion potential for the codon pairs that require only C•G-to-T•A and A•T-to-G•C editing, respectively. The C-to-T and A-to-G editing activities of Target-ACE were reduced by roughly up to half compared with those of Target-AID and ABE-7.10, respectively. Since the base editing spectra of Target-AID and ABE-7.10 were largely inherited in Target-ACE, most of the engineering strategies that have been adopted to optimize the effector domains of CBEs^23-26^ and ABEs^23^ could be additively adapted to improve the efficacy of Target-ACE without destroying its basic characteristics. While the overall low efficiency of Target-ACE is likely to be some structural unconformity of the large protein fusion of a total of 2,160 aa to express all of the three different functions originated from Cas9, PmCDA1 and TadA heterodimer, a protein engineering approach could also be applied to alleviate this effect or to produce a more optimized tool.

The aggregative bystander activities of two base editing effectors also confer the lower efficiency in precision source–estination codon conversions by producing more unexpected editing outcomes compared with those by independent CBEs or ABEs. This would be problematic, especially when unexpected base substitutions need to be minimized in the context of precision base editing. The bystander activity of Target-ACE could be reduced, using the engineered rAPOBEC1, APOBEC3A, PmCDA1, and TadA effector domains, which have narrow effective window sizes^13,27-30^. The use of a computational prediction method can also be considered to avoid genomic target regions that have high potential of being altered into unexpected sequences. While this study constructed simple conditional probability models for different base editing methods and achieved a certain level of performance in predicting base editing outcomes, other machine learning approaches could improve the prediction, particularly when coupled with base editing results for more target sites obtained by *en masse* DNA barcode-based mutational profile assays^16-18,31^ or by robotic automation.

Nevertheless, we here highlight that the multi-domain fusion strategy taken in Target-ACE is currently the best approach for inducing both C-to-T and A-to-G mutations in the DNA of the same cells. In general, the efficiency of gene delivery to cells is limited and decreased with increasing vector size. For gene therapy, retroviral and lentiviral transduction efficiencies are widely accepted to be maintained at a modest level for gene inserts of up to ∼10 kbp. Thus, as the insert sizes of Target-AID and ABE-7.10 are >6 kbp each including a promoter region, it would be difficult to transduce cells with a viral vector harboring both Target-AID and ABE-7.10. Even if this were successful, the heterologous base editing of infected cells would then suffer from the issue of activity competition of BE mix. Separate viral packaging followed by simultaneous or serial transduction of cells for the two different base editors would not be ideal for simultaneous C-to-T and A-to-G editing, especially when the infection efficiency is low, for example, in somatic tissues. The probability of multiple genes on different viral genomes being delivered to the same cell can be expected to exponentially decrease with the number of genes; this co-infection approach would therefore mostly produce undesired base editing outcomes by either of the base editors. Comparing these possible approaches with the existing CBEs and ABEs, Target-ACE of a total size of 7.1 kbp including a promoter region appears to be more beneficial for inducing heterologous mutations in the same cells.

Several studies have reported genome-wide off-target effects of CBEs^32,33^ and transcriptome-wide off-target RNA editing by some CBEs and ABEs, which have base editing effectors originating from RNA editing enzymes^30,34-36^. While the off-target RNA editing activity of Target-AID using the deoxycytidine deaminase PmCDA1 remains to be examined, Target-ACE is also expected to have at least a certain level of DNA off-target activity and carry over the RNA off-target effect from ABE-7.10. However, these undesired effects of CBEs and ABEs have been successfully attenuated by engineered mutations^30,34-38^. We expect that Target-ACE can incorporate these engineered mutations as well as other mutations mentioned above. Furthermore, other design considerations contributing to the optimization of wild-type Cas9^15,39-43^ should also be adaptable to Target-ACE, as seen in our previous example of Target-AID-NG, which is based on the engineered SpCas9-NG architecture that recognizes a single guanine PAM^15^.

Taking the obtained findings together, we have developed a new base editor Target-ACE by fusing two deaminase domains for C-to-T and A-to-G base editing and demonstrated its functionality expanding the patterns of sequence alteration at a targeted genomic position. More importantly, this paper presents the possibility of tethering two base editing effectors to a single genome-editing tool, shedding light on the potential for further diversifying base editing technologies with natural and engineered Cas9 variants.

## Acknowledgments

We thank members of the Yachie lab for useful discussions and critical assessment of this work. This study was mainly funded by the Uehara Memorial Foundation and partly supported by the New Energy and Industrial Technology Development Organization (NEDO), Japan Agency for Medical Research and Development (AMED) PRIME program (17gm6110007), Japan Science and Technology Agency (JST) PRESTO program (10814), the Naito Foundation, SECOM Science and Technology Foundation (all to N.Y.), and Japan Society for the Promotion of Science (JSPS) Grant-in-Aid for Scientific Research (16J06287) (to S.I.). S.I. was supported by a JSPS DC1 Fellowship; S.I., H.M., and N.M. were supported by a TTCK Fellowship; H.M. and N.M. were supported by the Mori Memorial Foundation; and N.M. was supported by the Yamagishi Student Project Support Program of Keio University.

## Author contributions

R.S., S.I., H.M., and N.Y. conceived and designed the study. R.S., S.I., and M. Tanaka constructed the plasmids. R.S., S.I., and N.M. performed the base editor assays and the library construction for high-throughput sequencing. S.I. established the base editor reporter cell lines. M.S. performed the fluorescence microscopy imaging. S.I. and H.M. performed the data analysis. M.S. and H.A. supported the high-throughput sequencing analysis. K.N., A.K., H.N., and O.N. supported the design of Target-ACE and provided materials. M. Tomita helped the computational analyses. R.S., S.I., H.M., and N.Y. wrote the manuscript.

## Online Methods

### Oligonucleotides

The oligonucleotides used in this study can be found in Supplementary Table 2.

### Plasmid construction

#### Base editor expression plasmids

All of the expression plasmids for base editors were standardized to the backbone used in pCMV-ABE7.10 (Addgene 102919). Two split fragments of the Target-AID-encoding region were amplified from pcDNA3.1_pCMV-nCas-PmCDA1-ugi pH1-gRNA(HPRT) (Addgene 79620) using primer sets RS045/HM129 and HM128/RS046 and assembled into the plasmid backbone amplified from pCMV-ABE7.10 using primers RS047/RS048 by Gibson Assembly, constructing pCMV-Target-AID (pRS0035). To construct pCMV-Target-ACE (pRS0045), the PmCDA1–UGI fragment was amplified from pcDNA3.1_pCMV-nCas-PmCDA1-ugi pH1-gRNA(HPRT) using primers RS051/RS046 and assembled into the plasmid backbone amplified from pCMV-ABE7.10 using primers RS047/RS052 by Gibson Assembly.

#### gRNA expression plasmids

gRNA spacer inserts were prepared by single pot reactions of phosphorylation and annealing of ssDNA pairs listed in Supplementary Table 2. Each T4 polynucleotide kinase (PNK) reaction sample was prepared for two ssDNA fragments in accordance with the manufacturer’s protocol (Takara) and set in a thermal cycler as follows: 37 °C for 30 min; 95 °C for 5 min; 70 cycles of 95 °C for 12 s and −1 °C/cycle; and then maintained at 25 °C. The annealed spacer inserts were then ligated into a pU6-gRNA cloning backbone (pSI-356) by Golden Gate Assembly using BsmBI (NEB) and T4 DNA ligase (NEB). The assembly was performed under the following thermal cycler conditions: 15 cycles of 37 °C for 5 min and 20 °C for 5 min; 55 °C for 30 min; and then maintained at 4 °C.

#### Lentiviral base editing reporter plasmids

To construct the lentiviral C-to-T base editing reporter plasmid (pLV-SI-112), the reporter cassette was amplified from pLV-eGFP (Addgene 36083) using 112-V4-BC2-FW/EGFP-BamH1-RV and cloned into EcoRI and BamHI sites of pLVSIN-CMV-Puro backbone vector (Takara) using T4 DNA ligase (NEB). The lentiviral A-to-G base editing reporter plasmid (pLV-SI-121) was constructed similarly, where the reporter cassette was amplified using EcoR1-ABE-T3-STOP-FW/EGFP-BamH1-RV instead.

The newly constructed plasmids were confirmed by Sanger sequencing. The plasmids used in this study can be found in Supplementary Table 1, the information on which has been deposited in Addgene.

### Selection of genomic gRNA target sites

In the selection of gRNA target sites, the regions with poly-cytosine repeats (poly-C), poly-adenine repeats (poly-A), alternating poly-adenine/cytosine repeats (poly-AC), or alternating poly-cytosine/adenine repeats (poly-CA) were selected from the human genome (hg19) as follows. In the initial screening, poly-C regions were required to have a poly-cytosine segment in a 7-bp sliding window for which the 5′ end shifted at intervals of 2 bp from −24 bp through −16 bp relative to PAM (five windows); poly-A regions were required to have a poly-adenine segment in a 6-bp sliding window for which the 5′ end shifted at intervals of 2 bp from −21 bp through −13 bp relative to PAM (five windows); and poly-AC and poly-CA regions were required to have the corresponding sequence pattern in a 6-bp sliding window for which the 5′ end shifted at intervals of 2 bp from −24 bp through −14 bp relative to PAM (six windows). We then eliminated the candidate regions that contained a homopolymer of ≥4 bp within a gRNA seed region spanning from −8 bp through −1 bp to PAM and those overlapping with an annotated exonic region to minimize the unexpected phenotypic effect. For every sliding window position of each of the repeat types, the top one or two candidate regions with the highest predicted gRNA activity scores^44^ were selected, and the final target regions with various sliding window positions for each repeat type were arbitrarily selected from ones for which PCR amplifications with candidate primer sets were successful. This resulted in the eight poly-C, eight poly-A, four poly-AC, and four poly-CA target regions. (Note that one of the poly-AC target regions was eliminated from the analysis due to the low sequence coverage.) All of the target regions with their corresponding gRNAs and amplicon sequencing primers can be found in Supplementary Table 2.

### Cell culture

HEK293Ta cells were purchased from GeneCopoeia and maintained in Dulbecco’s Modified Eagle’s Medium (DMEM) (Sigma) supplemented with 10% fetal bovine serum (FBS) (Thermo Fisher Scientific) and 1% penicillin–streptomycin (Sigma) at 37 °C with 5% CO_2_. Cells were routinely tested for mycoplasma contamination by nested PCR using culture medium as a template.

### Establishment of base editing reporter cell lines

For lentiviral packaging of the C-to-T and A-to-G base editing reporter plasmids, ∼3.0 × 10^5^ HEK293Ta cells/well were seeded in a six-well plate 1 day before transfection. In each packaging reaction, 489 ng of lentiviral plasmid was co-transfected with the two helper plasmids psPAX2 (Addgene 12260) and pMD2.G (Addgene 12259) of 366 ng and 122 ng, respectively, and 9.38 µL of polyethylenimine (PEI) (Polysciences) in 300 µL of phosphate-buffered solution (PBS). The next day, culture medium was changed to fresh medium, and 2 days later, culture supernatant containing lentiviral particles was harvested and aliquoted into 1.5 mL tubes. The viral sample was then stored at −80 °C before infection. For lentiviral infection, ∼2.0 × 10^6^ HEK293Ta cells/well were seeded on a six-well plate with 1 mL of cell culture medium, incubating for 24 h. The viral supernatant was then thawed at room temperature, mixed with 1 µL of 8 mg/mL polybrene (Sigma), and added to each cell sample. One day after infection, ∼5.0 × 10^3^ infected cells were re-seeded on a 96-well culture plate for functional titer measurement using CellTiter-Glo assay (Promega). Two days after infection, 2.0 µg/mL puromycin (Thermo Fisher Scientific) was added to the culture medium and incubated for 3 days to select the base editing reporter cells.

### Transfection assay

For the EGFP reporter assay of base editing, the established reporter cells were seeded in a 12-well plate at a density of ∼1.0 × 10^5^ cells/well. The next day, 2.7 µL of PEI, 100 µL of PBS, 600 ng of base editor expression plasmid reagent, and 300 ng of gRNA expression plasmid were mixed and incubated at room temperature for 15 min before application to each well for transfection. Microscope imaging was performed 3 days after transfection with InCellAnalyzer6000 (GE Healthcare) with a 20× objective lens. The BE mix plasmid reagent was prepared by mixing the Target-AID and ABE-7.10 plasmids at a 1:1 mass ratio. For base editing assay of the endogenous target regions, HEK293Ta cells were seeded in a 24-well plate at a density of ∼5.0 × 10^4^ cells/well. The next day, 1.2 µL of PEI, 50 µL of PBS, 300 ng of base editor expression plasmid, and 100 ng of gRNA expression plasmid were mixed and incubated at room temperature for 15 min before application to each well for transfection. All experiments were performed in triplicate.

### Library preparation for high-throughput sequencing

Three days after transfection, cell samples were transferred to a 96-well PCR plate and the culture medium was removed by aspiration. For cell lysis, 200 µL of 50 mM NaOH was added to each sample, heated at 95 °C for 10 min, and cooled down on ice, followed by the addition of 20 µL of 1 M Tris-HCl [pH 8.0]. Each target region was then PCR-amplified from the lysis sample as a template with the corresponding first HTS primer pair (Supplementary Table 2). Each PCR reaction was performed in a 20 µL volume including 1 µL of the template, 1 µL of 10 µM of each primer, 0.2 µL of Phusion polymerase, 5× Phusion HF buffer, and 1.6 µL of 2.5 mM dNTPs with the following thermal cycler conditions: 98 °C for 30 s; 30 cycles of 98 °C for 10 s, 60 °C for 10 s, and 72 °C for 10 s; followed by 72 °C for 5 min as a final extension. Each amplified product was further amplified in a volume of 20 µL containing 1 µL of 10-fold dilution of the first PCR product and custom Illumina index primers (Supplementary Table 2) with the following thermal cycler conditions: 98 °C for 30 s; then 15 cycles of 98 °C for 10 s, 65 °C for 10 s, and 72 °C for 10 s; followed by 72 °C for 5 min as a final extension. For each base editing method, the final PCR products of different target regions were electrophoresed in 2% agarose gels with 1× TBE buffer and the expected bands were pooled for extraction using FastGene Gel/PCR Extraction Kit (Nippon Genetics) to produce an Illumina sequencing library.

### Library quantification and sequencing

The Illumina sequencing libraries were quantified by qPCR using KAPA Library Quantification Kit Illumina (KAPA Biosystems) and multiplexed for deep sequencing. The final multiplexed library was again quantified by qPCR and sequenced with 30% PhiX control using Illumina MiSeq (Illumina MiSeq reagent v3 600 cycles for 2× 301 bp paired-end sequencing).

### High-throughput sequencing data analysis

The common adapter sequences were first mapped onto the amplicon sequencing reads using NCBI BLAST+ (version 2.7.0)^45^ with the blastn-short option, allowing identification of sample indices and demultiplexing of paired-end reads. The paired-end reads of each sample were then merged using FLASH (version 1.2.0)^46^ to generate single sequencing reads, which were further mapped onto the corresponding reference sequence of the target region using EMBOSS needle package (version 6.6.0)^47^ with the identity threshold of ≥80%. For each combination of base editor reagent and target site, the sequencing sample with the highest number of mapped reads was chosen from the triplicates for further analyses. The sequencing results for the EGFP transfection control were used to normalize sequencing errors.

### Base editing prediction framework

#### Training of base editing model

The amplicon sequencing results of different target regions were used to train a model for each base editor reagent to predict its editing outcomes and their frequencies for a given test sequence. To minimize the effects of potential sequencing errors in the training procedure, observed editing outcomes with relative frequencies of less than 1.0 × 10^−4^ are first eliminated from the dataset. Let *s*_*i*_ be the nucleotide base transition status at i bp position relative to PAM and *P* (*s*_*i*_) be the probability of *s*_*i*_. For each target region, *P* (*s*_*i*_) and *P* (*s*_*j*_ | *s*_*i*_) are calculated for every combination of *i* and *j* in a given area (*i* ≠ *j*). The base editing model is finally constructed as average *P*(*s*_*i*_) and average *P* (*s*_*j*_ | *s*_*i*_)across different training regions for which *s*_*i*_ is observed in ≥100 reads, represented by 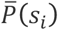 and 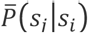, respectively.

#### Prediction of base editing outcomes

Let *S*_*m.n*_ be a base editing pattern in a window spanning from *m* bp through *n* bp relative to PAM, which can be alternatively represented by a string of transition statuses

*s*_*m*,_ *s*_*m* +1,_ …, *s*_*n* −1,_ *s*_*n*_. Using the training model, the frequency of a given outcome *S*_*m.n*_ in a test target region is predicted using the following equation:

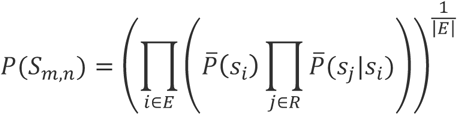

where *R* :={*x* ∈ *Z*|*m* ≤ *x* ≤ *n*}, *E* :={*x* ∈ *R*|s_x_ represents base transition}, 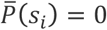 unless defined, and 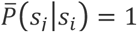 unless defined.

#### Validation of prediction framework

The prediction framework was evaluated by leave-one-out cross-validation for each of the base editor reagents. For every given test target region, frequencies of all of the base editing outcomes detected in the amplicon sequencing dataset for a window from −24 bp through −5 bp relative to PAM were predicted using the datasets of the other 22 target regions. The correlations between predicted and observed frequencies of the entire outcomes in all of the target regions were then evaluated by Pearson’s correlation coefficient.

#### Simulation of synthetic target sequences

To abstract the multi-dimensional base editing spectrum of each base editing method, we generated synthetic target regions harboring poly-C, poly-A, poly-AC, and poly-CA sequences spanning from −24 bp through −5 bp relative to PAM and the predicted frequencies of all of their possible outcomes with C-to-T and A-to-G base substitutions (2^20^ outcomes each). For each of the synthetic target regions, a base editing spectrum and di-nucleotide and tri-nucleotide co-editing score profiles were calculated from the predicted frequencies of all of the possible outcomes.

### Data availability

The high-throughput sequencing data are available at the Sequence Read Archive (PRJNA557370) of the NCBI.

