## Supplementary Materials for "A single CRISPR base editor to induce simultaneous C-to-T and A-to-G mutations"

Table of Contents


| | |
|---|---|
| **Supplementary Table 1** | List of plasmids used and generated in this study |
| **Supplementary Table 2** | List of oligonucleotides used in this study |
| **Supplementary Figure 1** | The EGFP signals obtained from the C-to-T and A-to-G base editing reporter cells for different base editing methods |
| **Supplementary Figure 2** | Base editing spectra of different methods for the poly-C regions |
| **Supplementary Figure 3** | Base editing spectra of different methods for the poly-A regions |
| **Supplementary Figure 4** | Base editing spectra of different methods for the poly-AC and poly-CA regions |
| **Supplementary Figure 5** | Comparison of C-to-T editing frequencies by different base editing methods |
| **Supplementary Figure 6** | Comparison of A-to-G editing frequencies by different base editing methods |
| **Supplementary Figure 7** | Co-editing spectra of Target-AID induced in the poly-C regions |
| **Supplementary Figure 8** | Co-editing spectra of ABE-7.10 induced in the poly-A regions |
| **Supplementary Figure 9** | Co-editing spectra of Target-AID in the poly-AC and poly-CA regions |
| **Supplementary Figure 10** | Co-editing spectra of ABE-7.10 in the poly-AC and poly-CA regions |
| **Supplementary Figure 11** | Co-editing spectra of BE mix in the poly-AC and poly-CA regions |
| **Supplementary Figure 12** | Co-editing spectra of Target-ACE in the poly-AC and poly-CA regions |
| **Supplementary Figure 13** | Average co-editing spectra of Target-AID and ABE-7.10 in the poly-AC and poly-CA regions |
| **Supplementary Figure 14** | Multi-base editing outcomes induced by Target-ACE and BE mix |
| **Supplementary Figure 15** | Performance of predicting relative editing outcomes |
| **Supplementary Figure 16** | Simulated co-editing spectra of different base editing methods for different synthetic target sequences |
| **Supplementary Figure 17** | Codon convertibility profiles for different base editing methods |

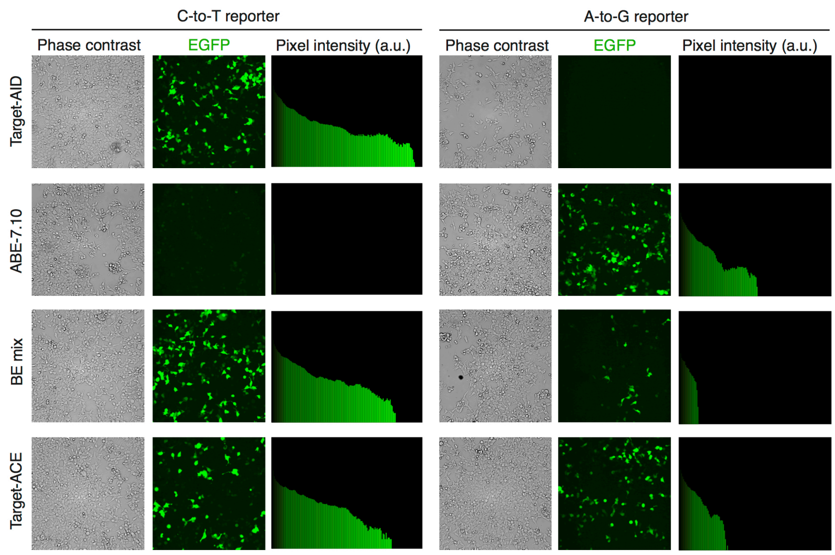


**Supplementary Figure 1.** The EGFP signals obtained from the C-to-T and A-to-G base editing reporter cells for different base editing methods. The pixel intensity distributions for the wider area images of Fig. 1d are shown.

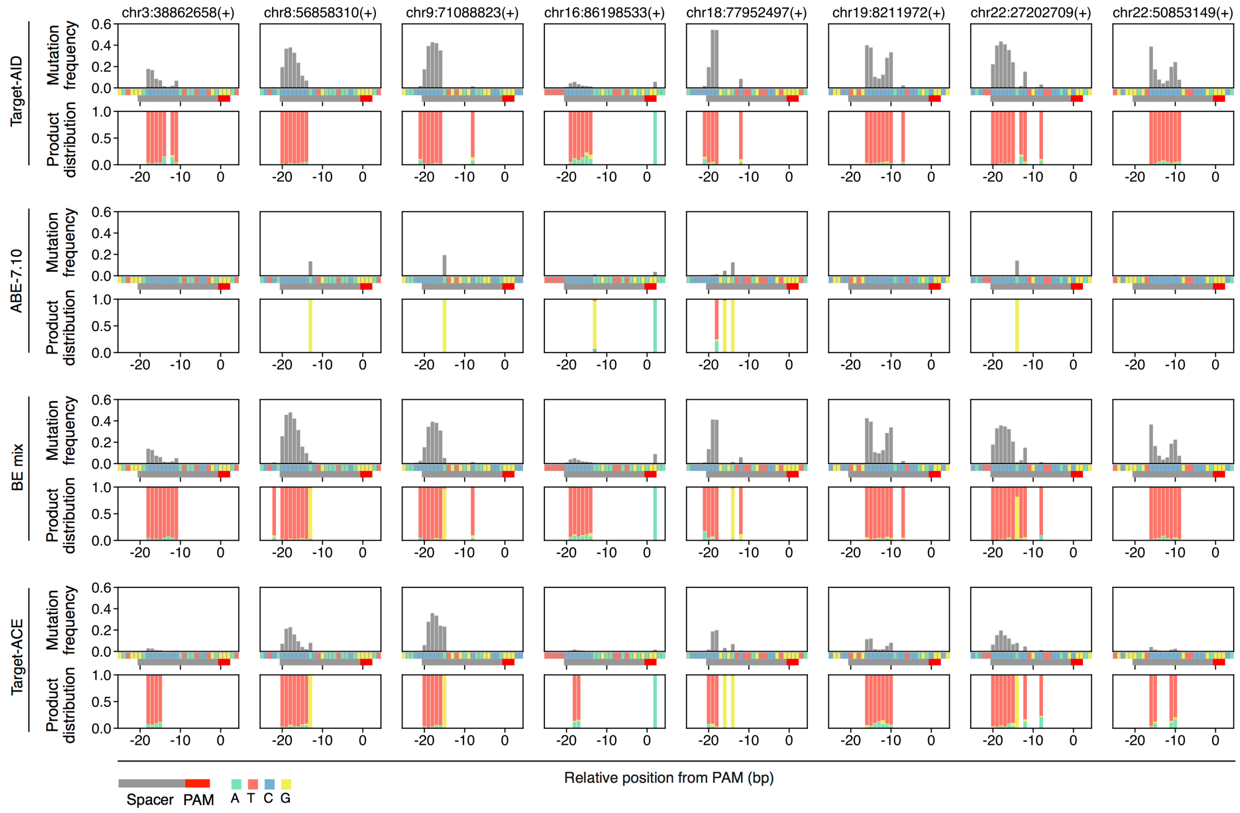


**Supplementary Figure 2.** Base editing spectra of different methods for the poly-C regions. Product distributions are shown for positions with base editing frequencies of ≥1%.

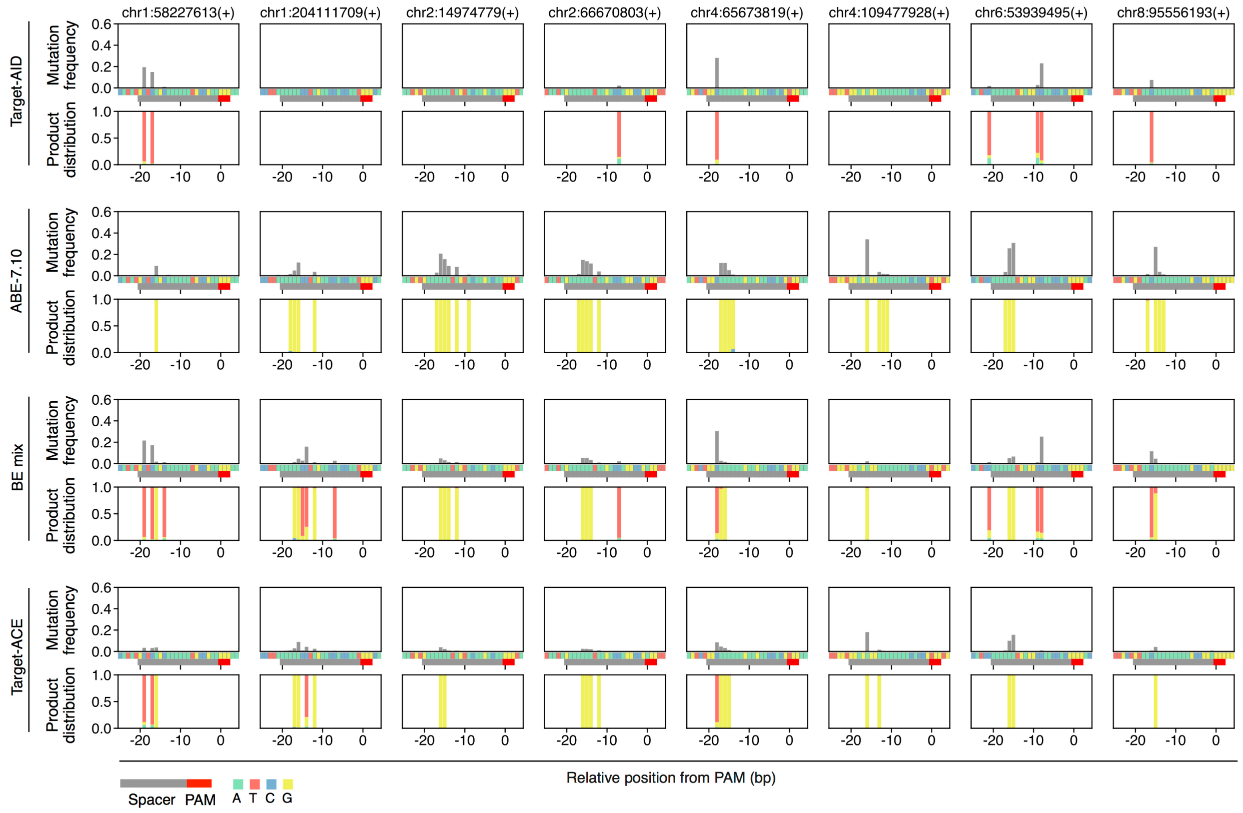


**Supplementary Figure 3.** Base editing spectra of different methods for the poly-A regions. Product distributions are shown for positions with base editing frequencies of ≥1%.

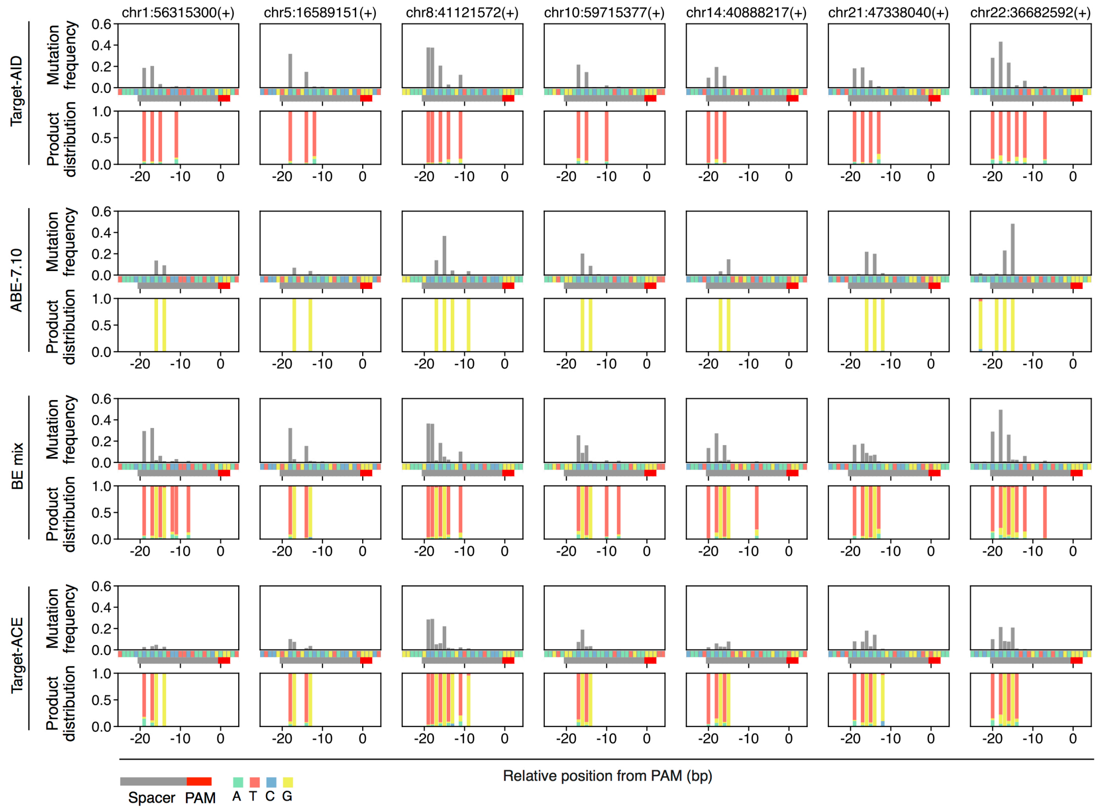


**Supplementary Figure 4.** Base editing spectra of different methods for the poly-AC and poly-CA regions. Product distributions are shown for positions with base editing frequencies of ≥1%.

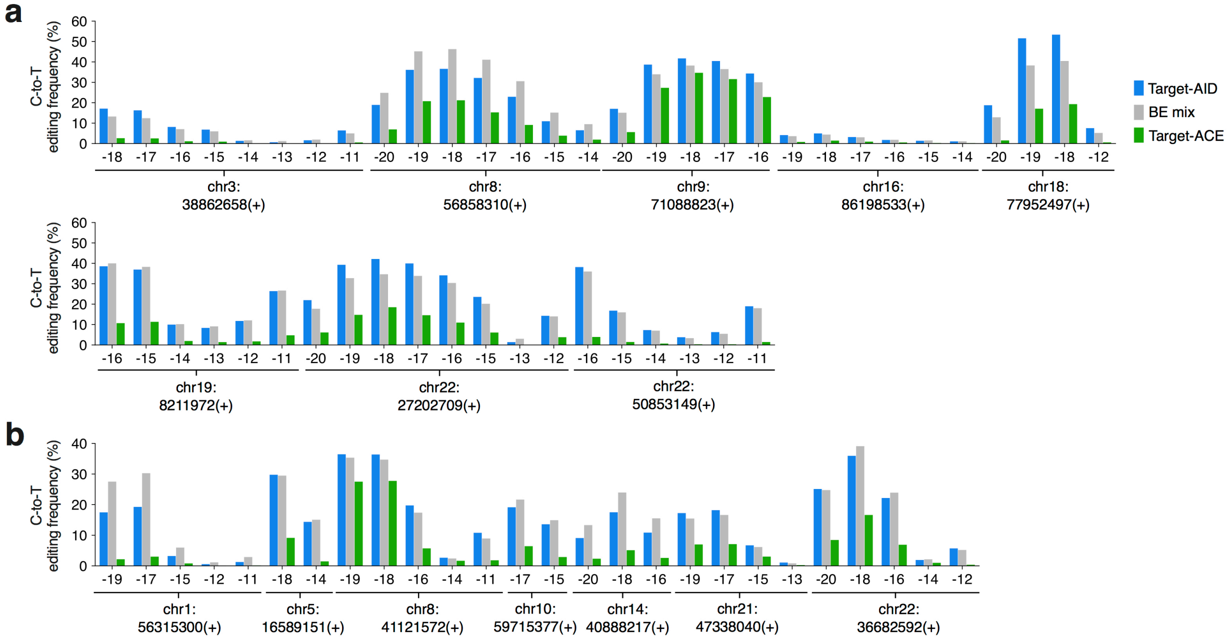


**Supplementary Figure 5.** Comparison of C-to-T editing frequencies by different base editing methods. (**a**) Poly-C regions. (**b**) Poly-AC and poly-CA regions. Each target region is denoted by its chromosome and the 5′ end position and strand direction of PAM.

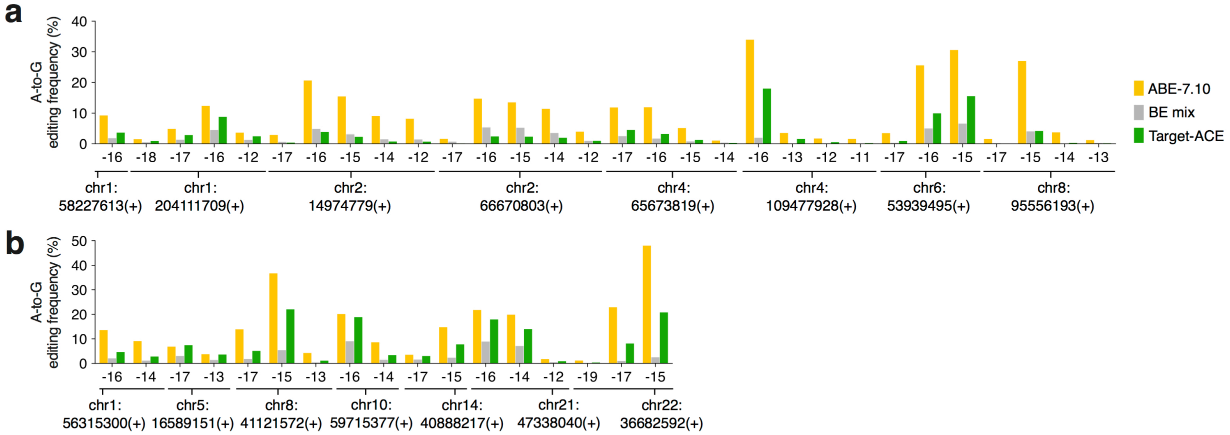


**Supplementary Figure 6.** Comparison of A-to-G editing frequencies by different base editing methods. (**a**) Poly-A regions. (**b**) Poly-AC and poly-CA regions. Each target region is denoted by its chromosome and the 5′ end position and strand direction of PAM.

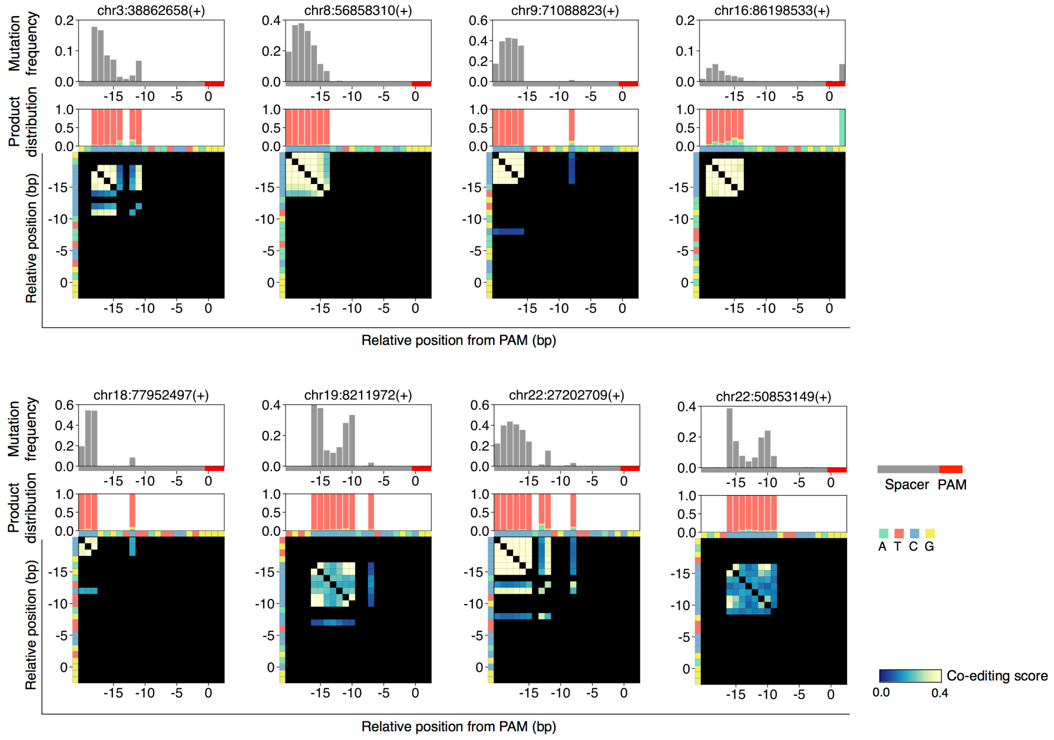


**Supplementary Figure 7.** Co-editing spectra of Target-AID induced in the poly-C regions. For each of the genomic poly-C regions, co-editing scores of all possible cytosine–cytosine combinations are shown along with their base editing frequencies and product distributions. Each target region is denoted by its chromosome and the 5′ end position and strand direction of PAM. Product distributions are shown for positions with base editing frequencies of ≥1%.

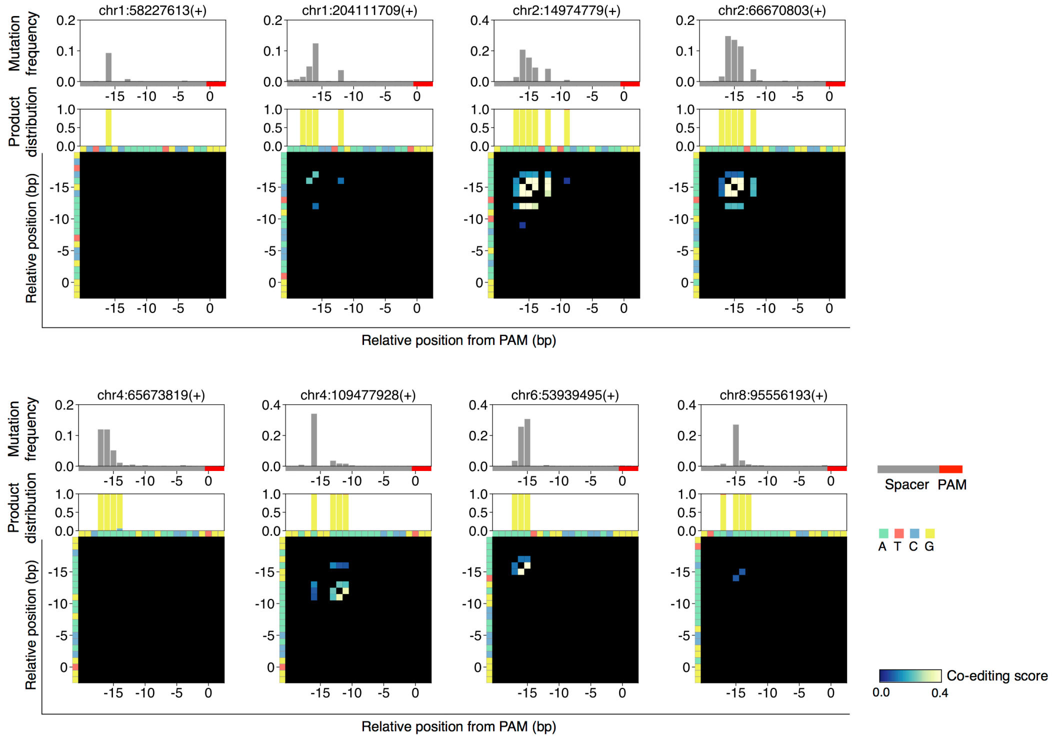


**Supplementary Figure 8.** Co-editing spectra of ABE-7.10 induced in the poly-A regions. For each of the genomic poly-A regions, co-editing scores of all possible adenine–adenine combinations are shown along with their base editing frequencies and product distributions. Each target region is denoted by its chromosome and the 5′ end position and strand direction of PAM. Product distributions are shown for positions with base editing frequencies of ≥1%.

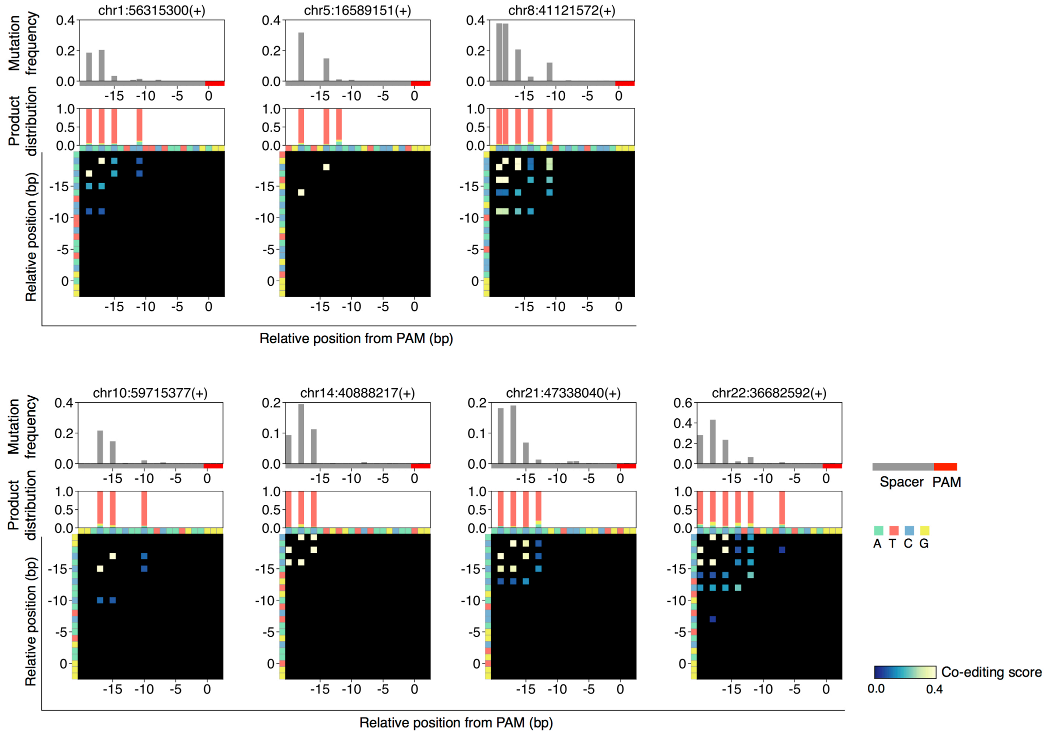


**Supplementary Figure 9.** Co-editing spectra of Target-AID in the poly-AC and poly-CA regions. For each of the genomic poly-AC and poly-CA regions, co-editing scores of all possible cytosine–cytosine, adenine–adenine, cytosine–adenine, and adenine–cytosine combinations are shown along with their base editing frequencies and product distributions. Each target region is denoted by its chromosome and the 5′ end position and strand direction of PAM. Product distributions are shown for positions with base editing frequencies of ≥1%.

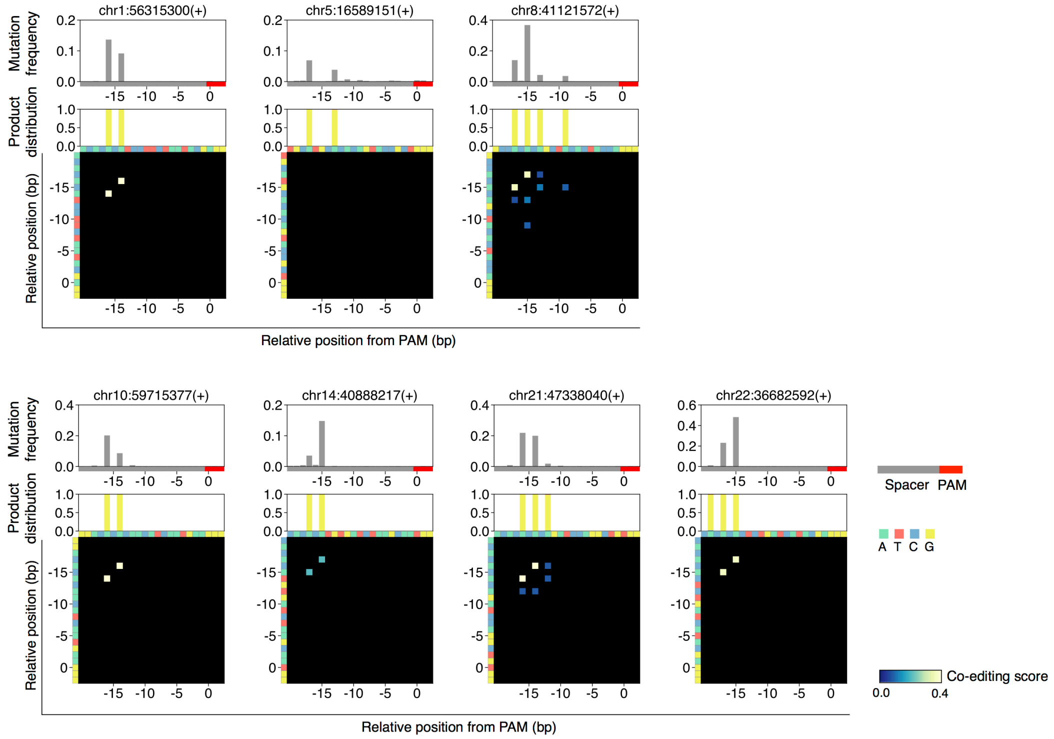


**Supplementary Figure 10.** Co-editing spectra of ABE-7.10 in the poly-AC and poly-CA regions. See the legends of Supplementary Fig. 9 for details.

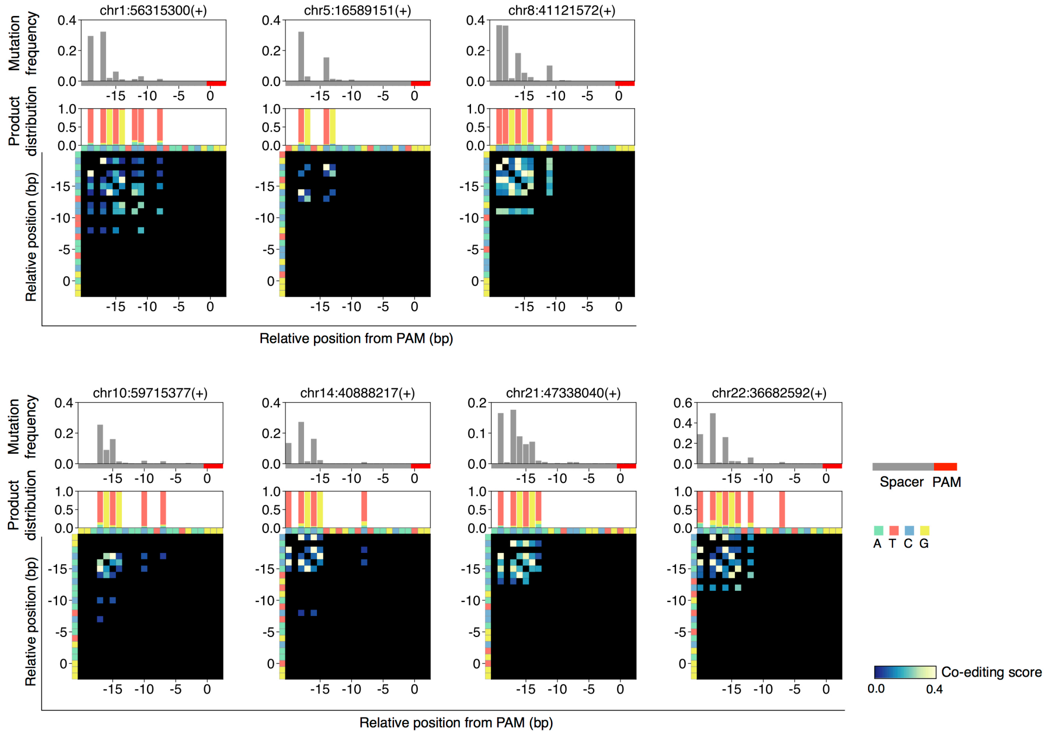


**Supplementary Figure 11.** Co-editing spectra of BE mix in the poly-AC and poly-CA regions. See the legend of Supplementary Fig. 9 for details.

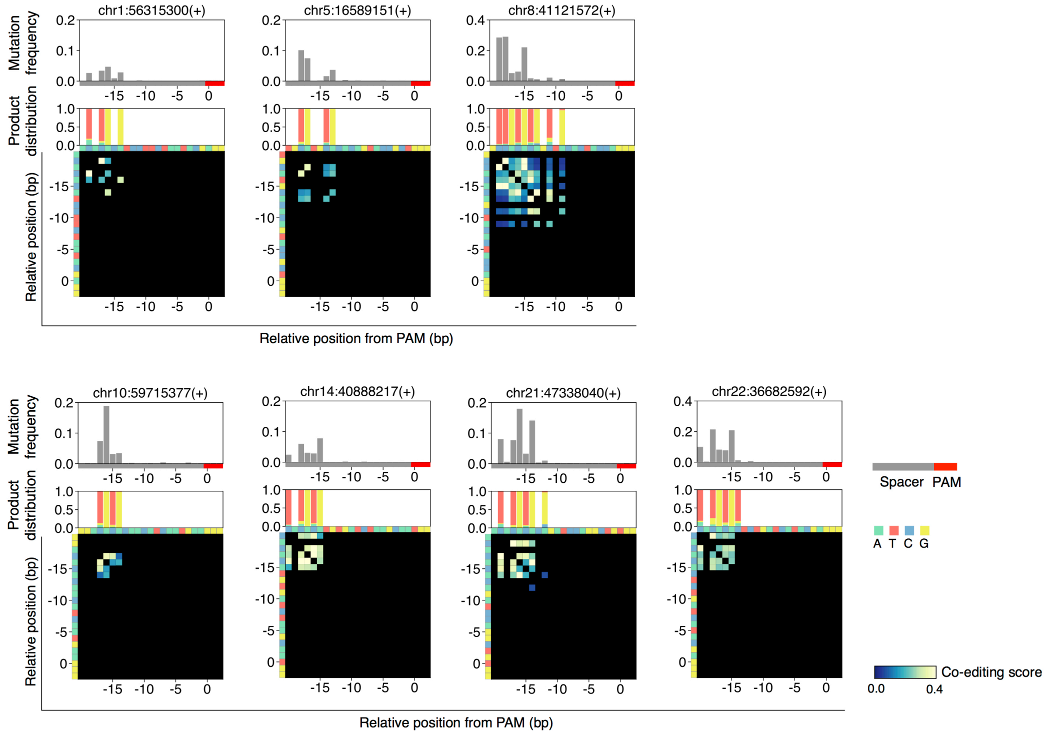


**Supplementary Figure 12**. Co-editing spectra of Target-ACE in the poly-AC and poly-CA regions. See the legend of Supplementary Fig. 9 for details.

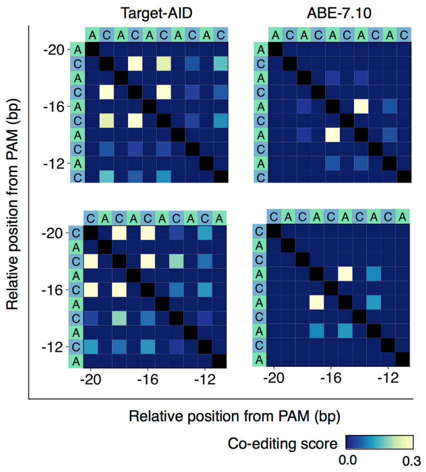


**Supplementary Figure 13.** Average co-editing spectra of Target-AID and ABE-7.10 in the poly-AC and poly-CA regions.

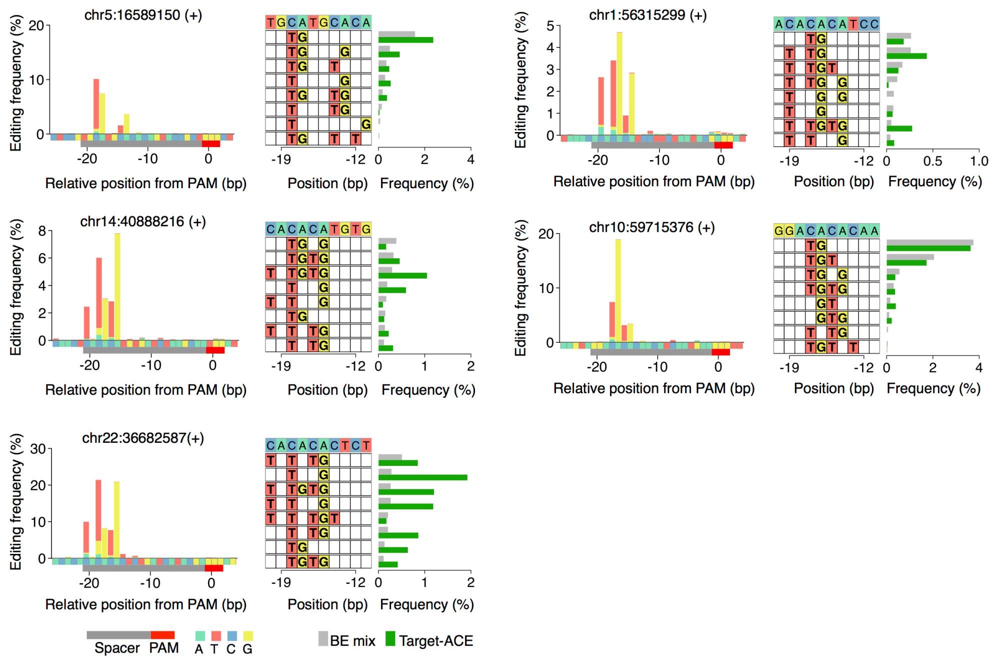


**Supplementary Figure 14.** Multi-base editing outcomes induced by Target-ACE and BE mix. Each target region is denoted by its chromosome and the 5′ end position and strand direction of PAM. In the left panel of each target region, base editing patterns and frequencies at different positions are represented by the color-coded x-axis for source nucleotide bases and the color-coded bars for destination nucleotide bases with their frequencies. In the right panel of each target region, the multi-base editing outcomes are shown for the top eight most common outcomes preferentially induced by BE mix.

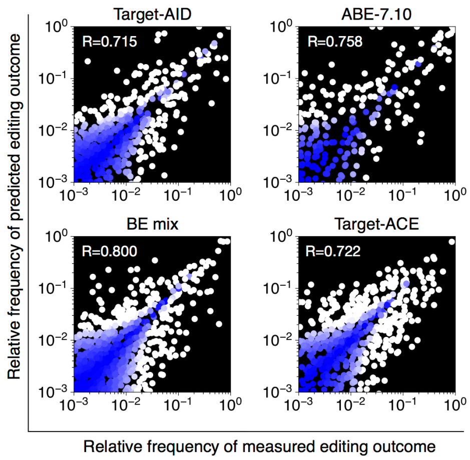


**Supplementary Figure 15.** Performance of predicting relative editing outcomes. The performance in predicting relative frequencies of different editing outcomes amongst all outcomes with base substitutions is shown for different base editing methods. The blue–white color scale indicates the Euclidean distance from the perfect prediction (diagonal).

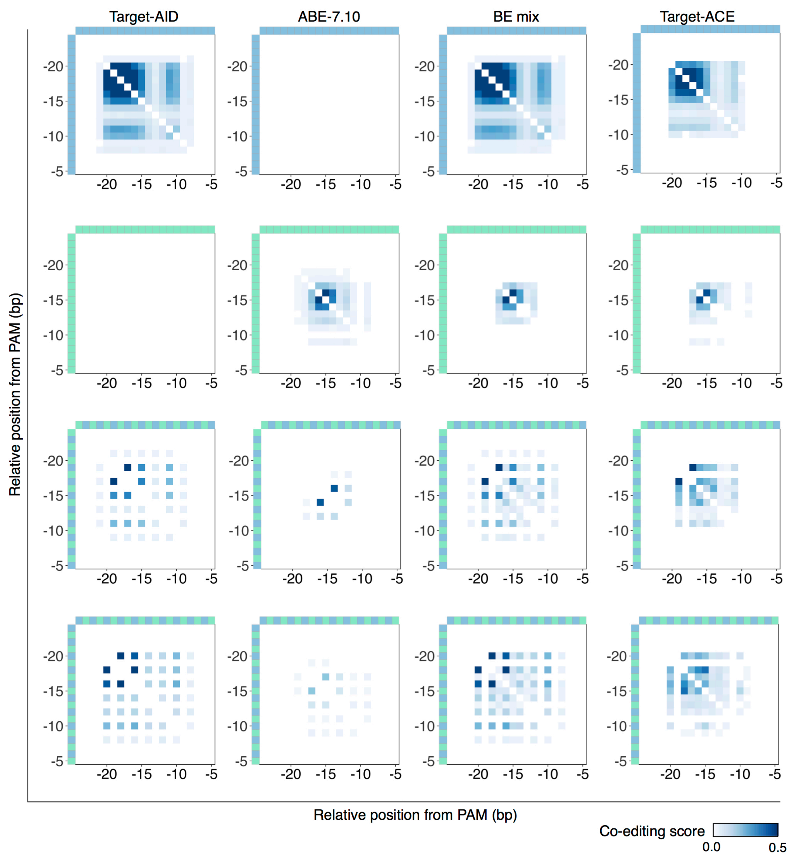


**Supplementary Figure 16.** Simulated co-editing spectra of different base editing methods for different synthetic target sequences.

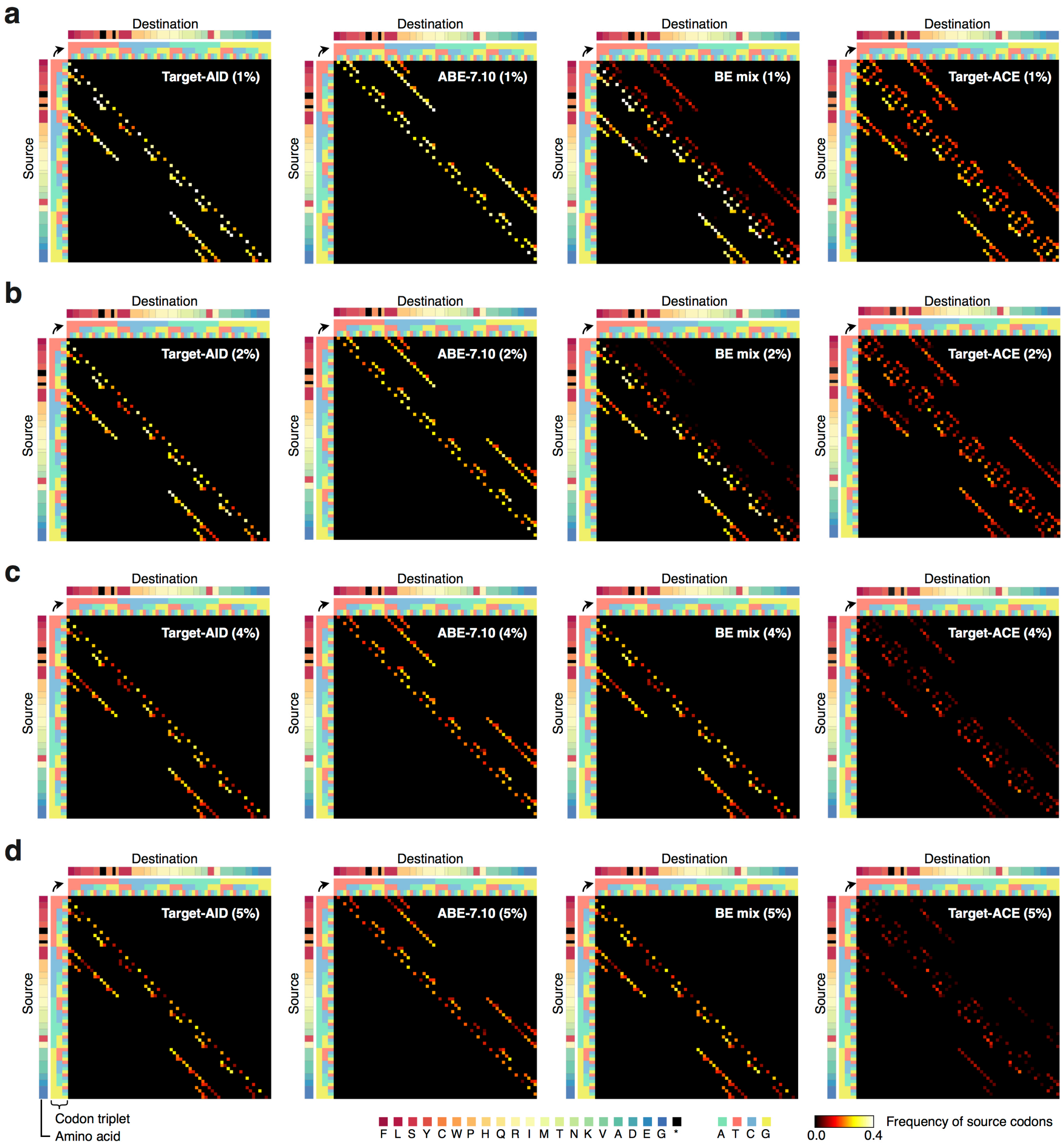


**Supplementary Figure 17.** Codon convertibility profiles for different base editing methods. For each source-to-destination codon conversion, the color scale indicates the fraction of the source codons in the human genome with predicted codon conversion potentials above a given threshold that keep surrounding nucleotide regions intact. The codon conversion potential thresholds of 1%, 2%, 4%, and 5% are presented in (**a–d**), respectively.
